# omicML: An Integrative Bioinformatics and Machine Learning Framework for Transcriptomic Biomarker Identification

**DOI:** 10.1101/2025.10.25.684517

**Authors:** Joy Prokash Debnath, Kabir Hossen, Md. Sayeam Khandaker, Shawon Majid, Md Mehrajul Islam, Siam Arefin, Preonath Chondrow Dev, Saifuddin Sarker, Tanvir Hossain

## Abstract

**Introduction:** Transcriptomic biomarker discovery has been a challenge due to variation in datasets and platforms, complexity in statistical and computational methods, integration of multiple programming languages, and intricacy of ML workflow to evaluate biomarkers. Standard workflows necessitate several stages (quality control, normalization, differential expression), typically executed in R or Python, resulting in bottlenecks for non-experts. Existing platforms have alleviated certain challenges by offering graphical interfaces for data loading, normalization, differential gene expression analysis, and functional analysis; nevertheless, they typically do not incorporate integrated machine learning procedures for biomarker selection.

**Method:** In this regard, we present omicML, an intuitive graphical user interface (GUI) that combines transcriptomic data analysis with machine learning (ML)-based classification via integrating R and Python packages/libraries. It supports both RNA-Seq and microarray data, automating preprocessing (data import, quality control, and normalization) and differential expression analysis. The tool annotates differentially expressed genes (DEGs) with descriptions, gene ontology, and pathway information and incorporates comparative analysis. Our extensive ML pipeline enables both supervised and unsupervised learning, integrates various datasets based on candidate gene signatures, standardizes and eliminates less significant features, benchmarks multiple ML classifiers with robust performance metrics (e.g., AUROC, AUPRC), assesses feature importance, develops single-gene and multi-gene predictive models, and systematically finalizes the biomarker algorithm. All functionalities are available in omicML, hence reducing the barrier for biologists without computational proficiency.

**Result:** In a case study, omicML identified a six-gene diagnostic model that distinguishes Mpox (monkeypox virus) infections from those caused by other viruses, including SARS-CoV-2, HIV, Ebola, and varicella-zoster. These results illustrate omicML’s capacity to discern clinically relevant biomarkers from complex transcriptome data.

**Conclusion:** Through the unified system, omicML (https://omicml.org), integrating data preprocessing, differential gene expression analysis, annotation, heatmap analysis, dataset integration, batch effect correction, machine learning approach, and functional analysis can diminish technical barriers and accelerates the conversion of expression data into diagnostic insights for clinicians and bench scientists.

## Introduction

Microarrays and RNA sequencing have been extensively utilized to investigate gene behavior in diseases, environmental stresses, or infections. These techniques generate mountains of data and translating high-dimensional expression data into robust biomarkers poses significant challenges, particularly for researchers without computational expertise. A biomarker discovery workflow demands proficiency in diverse tools for preprocessing, differential expression analysis, functional annotation, and advanced machine learning (ML), each requires specialized programming skills in R, Python, or Bash. Existing platforms streamline differential gene expression (DGE) analysis and pathway enrichment, but they typically lack ML pipelines for predictive biomarker discovery. To address these gaps, we present omicML, an interactive web application that integrates bioinformatics and machine learning workflows for the development of transcriptome biomarkers with clicks.

Over the years, numerous web-based platforms have been developed to optimize genomic and transcriptomic investigations, providing intuitive interfaces that enable complex data interrogation with minimal user burden. For example, iDEP (integrated Differential Expression and Pathway analysis) automates extensive gene ID conversion, provides comprehensive gene annotation, and integrates statistical and visualization techniques, including principal component analysis (PCA), DGE analysis, and pathway enrichment [1]. Similar platforms, such as START App (Shiny Transcriptome Analysis Resource Tool), Degust, and ShinyNGS, offer intuitive interfaces for clustering, PCA, expression visualization, and DGE analysis [2–4]. Furthermore, an open-source platform, Galaxy, provides a broad suite of web-based tools for genomic and differential analysis [5]. These platforms facilitate preliminary data exploration and DGE identification but do not inherently identify biomarkers using advanced ML. Additional specialized instruments also tackle many facets of gene expression analysis. IRIS-EDA facilitates DGE analysis through discovery-driven methodologies, including correlation analysis, heatmap creation, clustering, and PCA [6]. In contrast, Phantasus provides cross-platform datasets integration facilities from the Gene Expression Omnibus (GEO), allowing for data normalization and filtering before differential analysis [7]. The GEO2R tool in the NCBI, GEO portal, enables pairwise comparisons of expression data; however, it depends on the pre-normalized data provided by submitters and does not execute batch correction or standardize microarray and RNA-Seq processing. GEO2R excludes any gene that displays at least one NA value in any of the samples in the comparison [8].

Despite the advancements in transcriptome analysis provided by the aforementioned technologies, none are specifically engineered for predictive biomarker modeling utilizing comprehensive ML methodologies. To address this significant deficiency, we offer a universal algorithmic pipeline capable of predicting biomarkers linked to various diseases, environmental pressures, and other situations across several species, based on the differentially expressed genes revealed in preliminary research. omicML extends existing capabilities with the following features:(1) automated preprocessing of input data (imputation of missing values, batch effect correction), (2) compatibility with both RNA-Seq and microarray datasets, enabling cross-platform analysis, (3) annotation support for 367 species, providing broad taxonomic coverage, (4) integrated ML pipelines for data standardization, feature selection, benchmarking, nested coss-validation, hyperparameter tuning, feature importance, single-gene model building, multi-gene model (biomarker algorithm) building, model finalization, and (5) embedded network analysis and functional enrichment to contextualize candidate biomarkers within biological pathways. By integrating these processes, omicML distinctly connects transcriptomic analysis with predictive biomarker modeling, enabling users to convert DEG lists into actionable diagnostic or therapeutic targets without requiring coding expertise. omicML has been designed using six general modules **(Figure 01)**.

**Figure 1.**
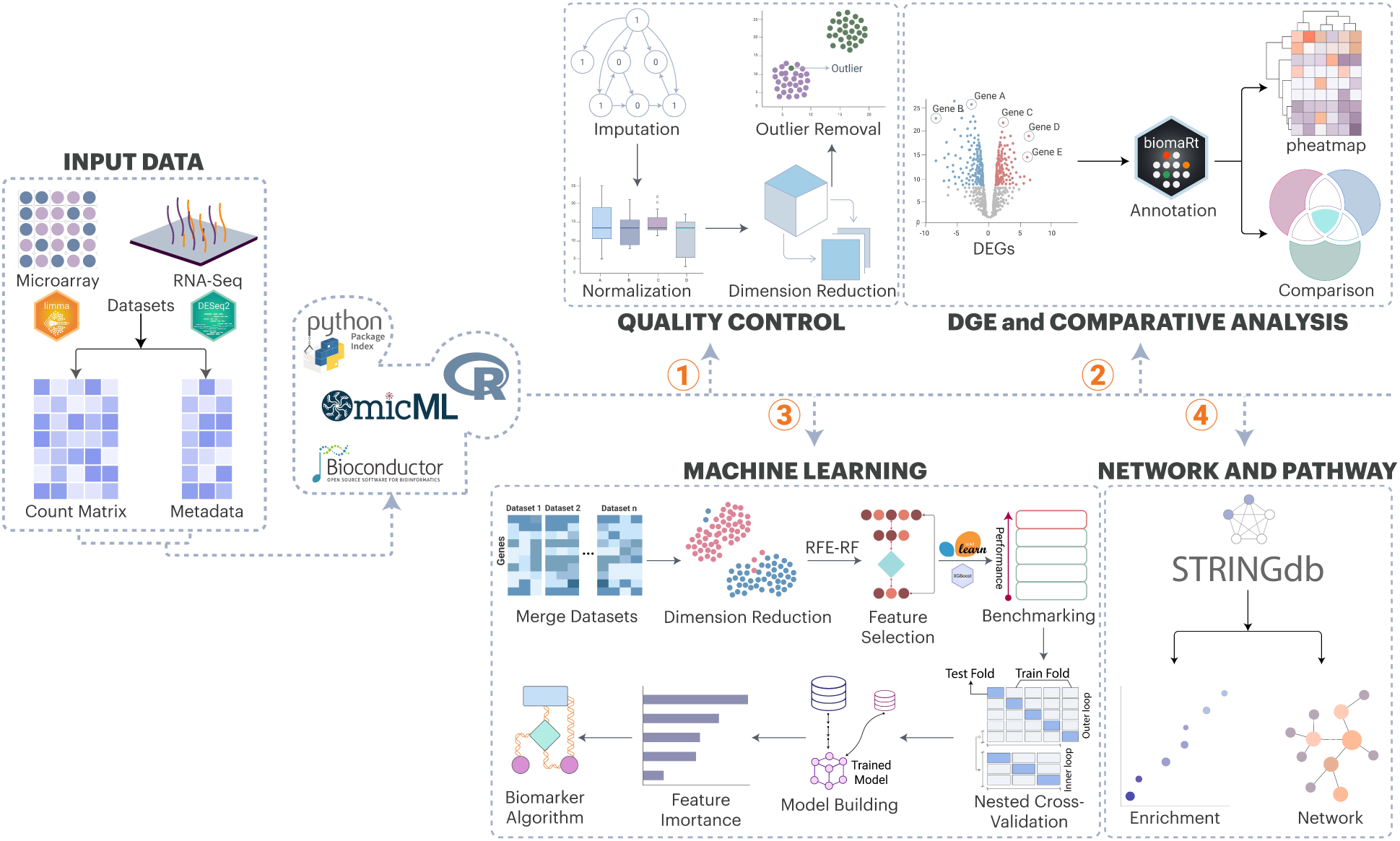
Overview of omicML and its modules.

This paper also illustrates the effectiveness of omicML through a case study on monkeypox virus (MPXV) infections. In the light of the identification of a novel clade of 2022 Mpox and their biomarkers in our previous study [9], we have now used heterogeneous transcriptomic datasets to identify the conserved biomarkers across different clades and cell models. omicML identified 34 shared DEGs through comparative analysis and employed a machine learning pipeline to prioritize six high-confidence biomarkers. Among these, RRAD emerged as the most robust single-gene predictor of MPXV infection. This example highlights omicML’s capacity to democratize ML-driven biomarker discovery, providing a reproducible, scalable, cross-platform workflow for users navigating complex transcriptomic data.

## Materials and Methods

Key steps are summarized in **Figure 2** which outlines the omicML workflow within a comprehensive bioinformatics pipeline for transcriptomic data analysis, focusing on biomarker discovery and functional interpretation.

**Figure 2.**
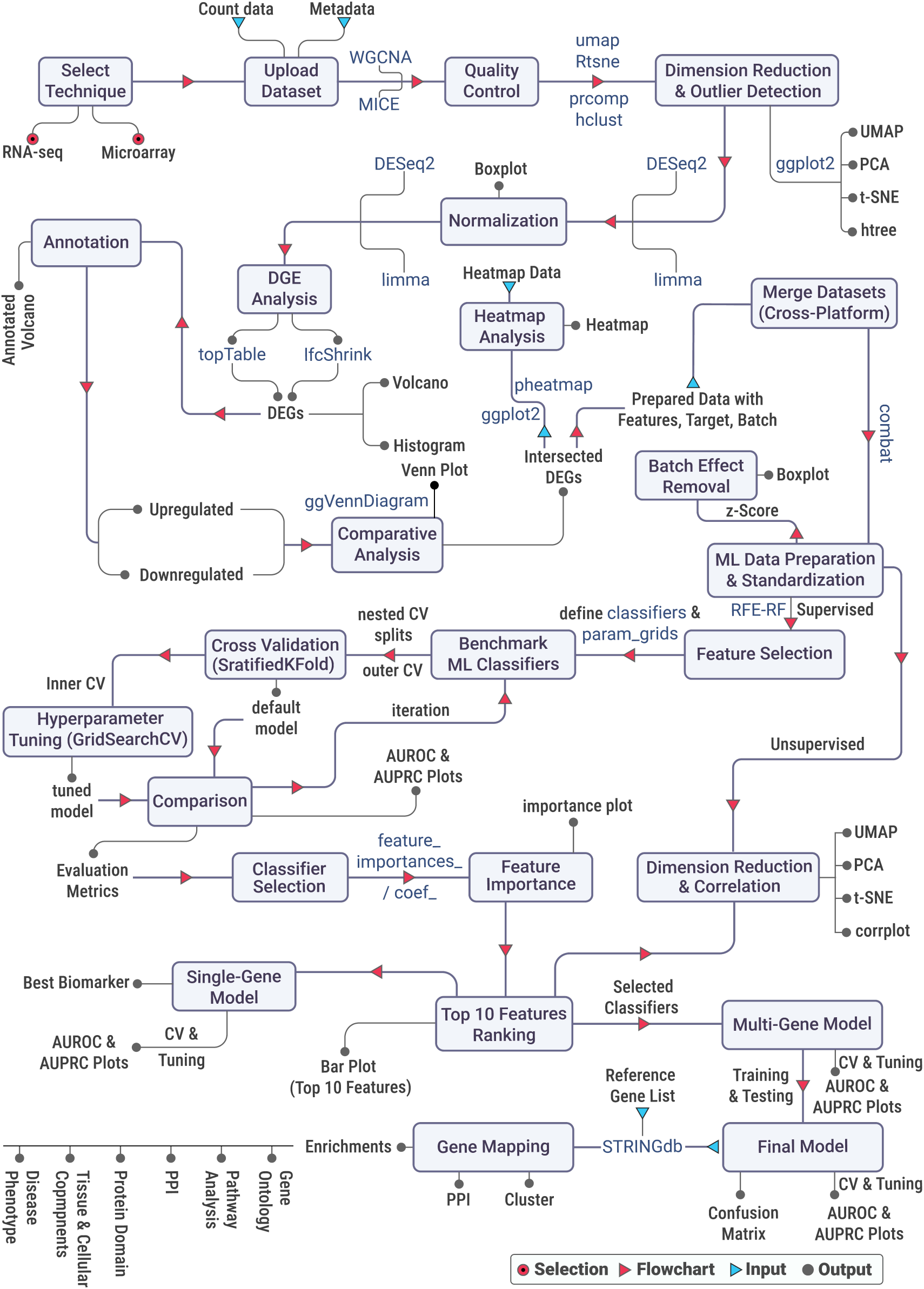
Detailed workflow of omicML

**Figure 3.**
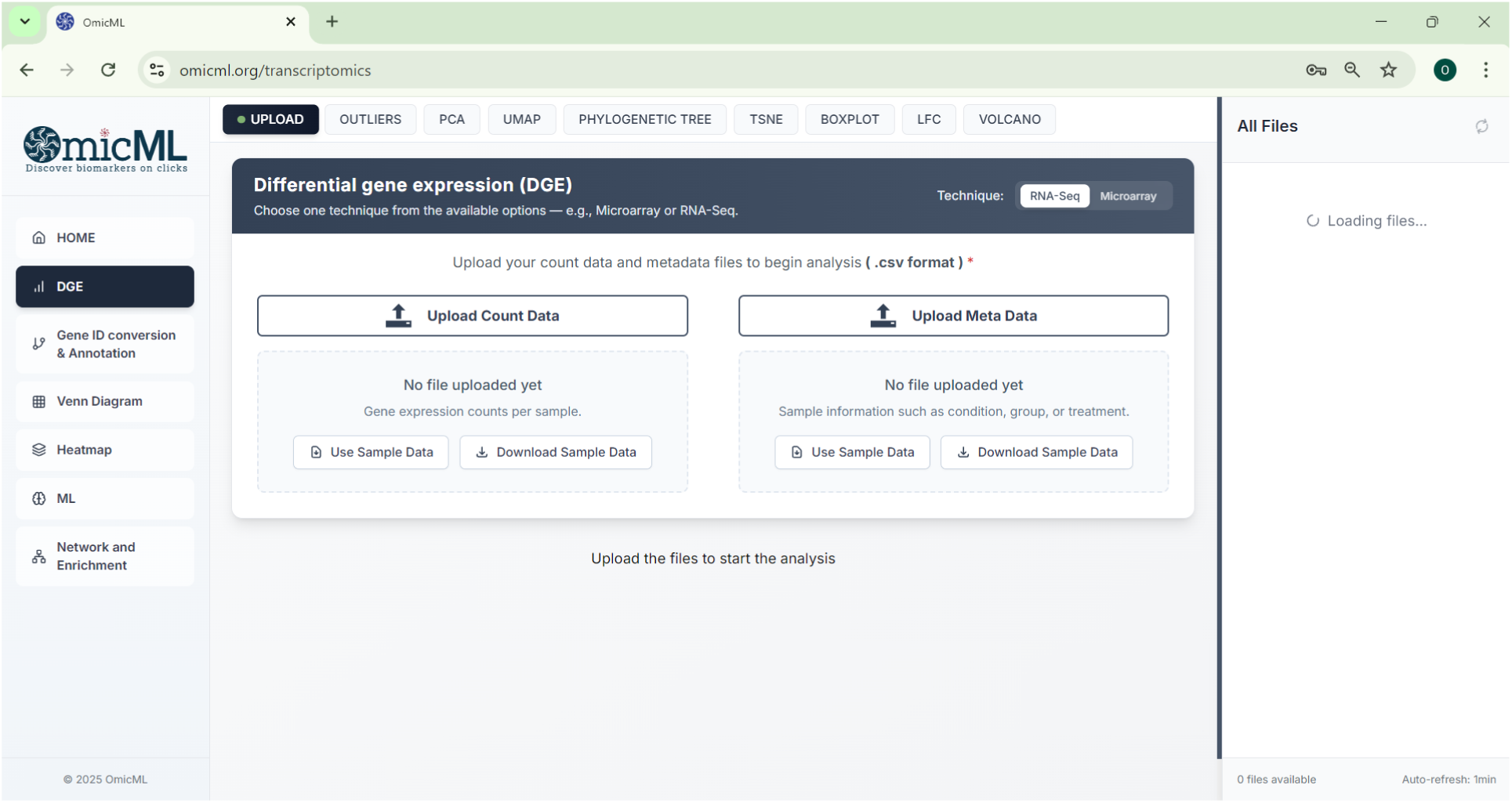
The snapshot represents the interface of the omicML web tool

### Data Acquisition and Input

The analytical pipeline within the omicML framework commences with the acquisition of primary input datasets of raw sequencing derived from either microarray or RNA-Seq experiments. Initial data can either be generated *de novo* via web-based analytical suites (e.g., Galaxy [5]) or retrieved from curated repositories such as Gene Expression Omnibus (GEO) [10], ArrayExpress [11], The Cancer Genome Atlas (TCGA) [12], the Sequence Read Archive [13].The structured input system—an expression matrix and a metadata is designed to minimize errors during processing and provide a robust foundation for ML-driven biomarkers discovery.

### Preprocessing, Dimensionality Reduction, and Outlier Detection

To ensure data integrity, initial preprocessing is employed by the filtration through WGCNA [14] and the imputation technique to mitigate low-variance genes and missing values respectively. Subsequently, dimension reduction techniques are used to visualize the high dimensional and complex gene expression data into lower dimension spaces. Additionally, hierarchical clustering (dendrogram-based view) is conducted to explore the grouping of samples facilitating the detection of potential outliers (user exclusion if necessary).

### Normalization and Batch Correction

Normalization corrects the systematic biases and “uninteresting” factors, ensuring that observed differences between experimental conditions accurately reflect true biological variation. For microarray data, quantile normalization is applied for harmonizing intensity distributions across arrays. RNA-Seq data is normalized in median-of-ratios approach.

### Differential Gene Expression (DGE) Analysis

In DGE analysis, differentially expressed genes (DEGs) are pointed out through pairwise comparison between samples with two different types of conditions. DGE analysis is employed through platform-specific statistical frameworks. DEGs are defined by |LFC| > 1 and padj < 0.05, visualized via volcano plots annotated with gene IDs.

### Gene ID Conversion and Annotation

Gene (DEGs) IDs (Ensembl and Entrez) resulted in DGE analysis are converted to gene symbol and annotated with description using the R package the *biomaRt* [15]—incorporates gene annotation data for 367 organisms (214 from Ensembl and 153 from Ensembl Plants). Gene ID conversion ensures the visualization of volcano plots with respective gene symbols (if available) instead of gene IDs (Ensembl or Entrez).

### Comparative Transcriptomics and Integrative Analysis

Comparative analysis of the outcomes in DGE analysis enhances our understanding of shared and unique gene expression patterns, revealing condition-specific molecular signatures. The result is visualized in a venn diagram using the *ggVennDiagram* [16] package, which can efficiently handle up to 7 gene sets. For analyses involving more than 7 gene lists, the results are visualized using an upset plot, offering a clear and scalable representation of complex gene set intersections. Users can analyze one or multiple datasets and compare the DEGs to identify intersecting genes shared across different contrasts as well as unique gene sets specific to individual contrasts. Log-fold metrics of both microarray and RNA-seq are integrated to generate the data for heatmap analysis.

### Machine Learning Analysis

A suite of advanced ML frameworks was leveraged to systematically evaluate the classification efficacy of the DEGs as candidate biomarkers derived from transcriptomic profiles. The predictive performance of the DEGs was quantified through iterative feature optimization and cross-validation, identifying genes with robust discriminatory power. This integrative pipeline bridges transcriptomic discovery and computational validation, thereby ensuring the robustness of putative biomarkers in complex biological matrices.

### Data Preprocessing, Merging, and Batch-Effect Correction

Raw multi-platform datasets are subjected to a systematic preprocessing workflow encompassing variance stabilization, and global normalization to attenuate platform-derived technical variability and facilitate cross-study comparability. Heterogeneous datasets (e.g., RNA-Seq and microarray) were integrated into a unified expression matrix, annotated with metadata columns for conditions and batch identifiers. Platform-induced technical artifacts are corrected via the *ComBat* [17], followed by Z-score to standardize feature distributions prior to downstream analyses.

### Dimensionality Reduction and Unsupervised Analysis

Unsupervised learning techniques, including Principal Component Analysis (PCA), t-distributed Stochastic Neighbor Embedding (t-SNE), and Uniform Manifold Approximation and Projection (UMAP), are applied to the normalized dataset to reduce dimensionality and reveal intrinsic data structures. Sample-to-sample relationships are further assessed by computing a Pearson correlation matrix, visualized as a correlation heatmap.

### Feature Selection

Feature selection is performed using the Recursive Feature Elimination (RFE) algorithm [18] with a Random Forest classifier as the base estimator. In this analysis, the top 50% of are retained by default. Finally, the dataset is reduced to retain only the selected features for subsequent modeling and analysis.

### Model Selection, Nested Cross-Validation, Hyperparameter Tuning and Model Benchmarking

Six ML classifiers, linear and tree-based methods, including Logistic Regression (LR), Extra Trees (ET), Random Forest (RF), XGBoost (XGB), Gradient Boosting (GB), and AdaBoost (AB), are selected for benchmarking experiment using Python libraries *scikit-learn* [19] and *XGBoost [20]*. To evaluate model performance and extract optimal hyperparameters, two-tiered nested cross-validation (CV) [21] is implemented. The 5-fold outer CV is used for model evaluation to maintain original class balance. In each of the five iterations, 4-folds (80% of the data) are utilized for training and hyperparameter selection; the remaining 1-fold (20%) is employed for testing. This process is repeated in all the 5-folds. To further optimize the model’s performance, hyperparameter tuning is conducted. Within each outer training set, an inner 3-fold Stratified CV (2-fold for training and 1-fold for testing) is performed for hyperparameter tuning, evaluating all combinations of hyperparameter using *GridSearchCV [20]*. A baseline evaluation is also performed using each classifier’s default hyperparameters under the same outer CV framework. For every held-out test fold, several metrics—including Accuracy (ACC), balanced accuracy (BACC), precision (PREC), recall (REC), F1 score (F1), AUROC, area under the precision-recall curve (AUPRC), Matthews correlation coefficient (MCC), Cohen’s kappa (KAPPA), and log loss (LOGLOSS)—are computed for both the default and tuned pipelines.

### Performance evaluation

The classification performance of each model is assessed through the calculation of the AUROC and AUPRC. By default, the model with the highest AUPRC and AUROC scores is selected for further analysis. But users can select any of the models which aligns well with their study.

### Feature Importance

For the selected model, feature importance is evaluated by calculating the mean decrease in accuracy, which measures the decrement of a model’s accuracy for the removal of individual feature. Based on this metric, the top 10 features are ranked, reflecting the features that contributed the most to the model’s accuracy in classifying samples.

### Single-Gene Model Building to Determine the Best Biomarker

For each of the top 10 features, a separate single-gene model is trained, and cross-validated using nested CV. The evaluation metrics (e.g., AUROC, AUPRC, ACC) are calculated accordingly. AUROC value indicates the ability of a feature (biomarker) to distinguish between, while the AUPRC value detects the ability of a feature to detect true positive case of the target condition. The single-gene model with the highest AUPRC and AUROC is considered as the best biomarker for the contrast.

### Feature Ablation Study to Finalize Multi-Gene Model (Biomarker Algorithm)

To finalize the multi gene-model, the top ten features are incrementally reduced by removing the least important feature one at a time. The multi-gene model yielding the best AUPRC along with other matrices is selected as the prediction model. Nested CV, hyperparameter tuning, and model evaluation (e.g., AUROC, AUPRC, ACC) are performed to finalize the multi-gene model (biomarker algorithm).

### Network and Functional Enrichment Framework

A set of genes’ symbols are uploaded and selected their corresponding organism to identify the PPI network and other enrichments. Initially, PPI networks are constructed using ***STRING-db*** [22] for the input gene symbols. Consequently, the common genes between the neighbor genes and query genes (need to be uploaded by user) are selected and their interaction network is visualized. The common genes are then analyzed to obtain their enrichments including Component, Process, Function, WikiPathways, KEGG, Reactome, HPO, Diseases, Pfam, SMART, InterPro, TISSUES, Compartments, NetworkNeighbors, PMID, Keyword with FDR<0.05. The top enriched terms are visualized using ***ggplot2***. Moreover, users can identify neighbor genes and upload multiple genes directly to obtain their interaction networks and enrichments.

### Backend and Frontend Development

The analytical pipelines are developed as APIs using *FastAPI [23]* to handle HTTP requests and enable communication with Python. R-based computations are executed using the *rpy2 (*https://rpy2.github.io/) interface. This architecture enables seamless interoperability between Python and R. The entire workflow is containerized using *docker [24]* to configure and manage the server. The frontend is developed using *Next.js (*https://nextjs.org/) and *TypeScript (*https://www.typescriptlang.org/) to create an interactive Graphical User Interface (GUI). *Next.js* is chosen for its server-side rendering (SSR) capabilities, improved SEO, and efficient static site generation (SSG). *TypeScript* is incorporated to enhance code maintainability and reliability. To Deploy the full server, EC2 (https://aws.amazon.com/ec2/) service of AWS is utilized.

## Results

We developed omicML, an intuitive web application integrating R and Python libraries to streamline transcriptomic analysis and biomarker prediction through a six-phase workflow: (i) differential gene expression analysis, (ii) gene annotation, (iii) comparative analysis, (iv) heatmap analysis, (v) learning analysis and validation, and (vi) functional analysis. The platform automates preprocessing and employs dimensionality reduction alongside incorporates exploratory tools (histograms, volcano plots) to identify DEGs. Machine learning workflows rigorously prioritize and validate biomarkers via data standardization, feature selection, benchmarking, nested cross-validation, hyperparameter tuning, feature importance, single-gene model building, multi-gene model (biomarker algorithm) building, model finalization, culminating in predictive models selected by evaluation metrics (e.g., AUROC, AUPRC, accuracy). By uniting statistical, ML, and functional analyses within an intuitive graphical user interface, omicML enables researchers without rigorous programming expertise to rapidly translate expression data into validated biomarkers and novel biological hypotheses with some clicks only.

## Case Study

### markerMPXV: Upregulation of RRAD and Building of a Biomarker Algorithm for Mpox Virus Infection

To demonstrate the potential of omicML, we applied it to analyze real biological data, selected from Gene Expression Omnibus (GEO), NCBI, of human cell models infected with the monkeypox virus (MPXV) strains. The dataset, GSE11234 [25], comprises gene expression data generated via expression profiling by array experiment of MPXV (Zaire strain)-infected dermal fibroblasts and monocytes. The other dataset, GSE219036 [26], employs high-throughput sequencing of transcriptomes of human keratinocytes infected with three distinct MPXV strains: Clade I (historically endemic), Clade IIa (prior endemic), and Clade IIb (exclusively classified during the 2022 global outbreak). Clade IIb shows distinguished expression pattern than the other clades, we identified in our previous article [9].

#### Pre-processing, and Normalization of Datasets

In the microarray dataset, data of fibroblast (16 samples) and monocyte (36 samples) cell lines were normalized to mitigate technical variability and ensure cross-sample comparability, with boxplots (**Figure 4A-D**) visualizing the reduction in technical biases and improved post-normalization distribution consistency. For the RNA-Seq dataset, expression distributions before and after normalization of keratinocyte cell-line samples were visualized in boxplots (**Figure 4E-F**)

**Figure 4.**
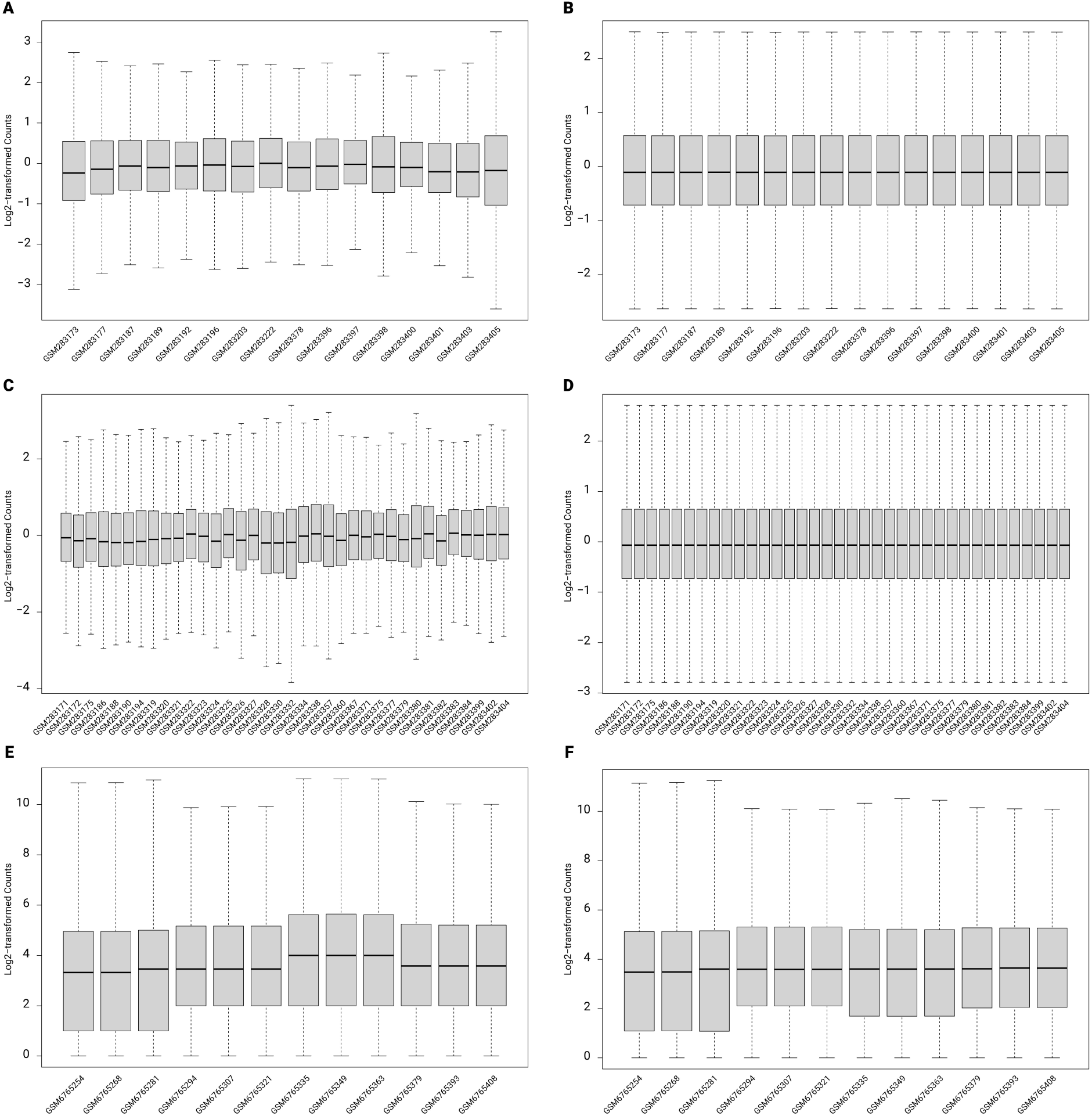
Expression pattern of datasets before and after normalization. The distribution of expression data in the fibroblast (A-B), monocyte (C-D), and keratinocyte (E-F) cell lines are shown. Each of the boxes represents 50 percent (median) of the data. The lower boundary (Q1) marks the 25th percentile, and the upper boundary (Q3) denotes the 75th percentile, with the central line indicating the median. 50% of data points lie above the median, and 50% fall below it, offering a clear statistical summary of gene expression variability within each cell type and normalization state.

UMAP **(Figure 5A, C, E)** and hierarchical clustering (**Figure 5B, D, F**) analyses revealed distinct expression patterns between MPXV-infected and mock cell types. Hierarchical clustering identified four and two MPXV-infected samples in fibroblast (**Figure 5B**) and monocytes (**Figure 5D**) respectively as outliers, underscoring technical or biological variability. Likewise, keratinocyte cell-lines (**Figure 5E-F**) exhibited high homogeneity, tight clustering with no misclassification between groups.

**Figure 5.**
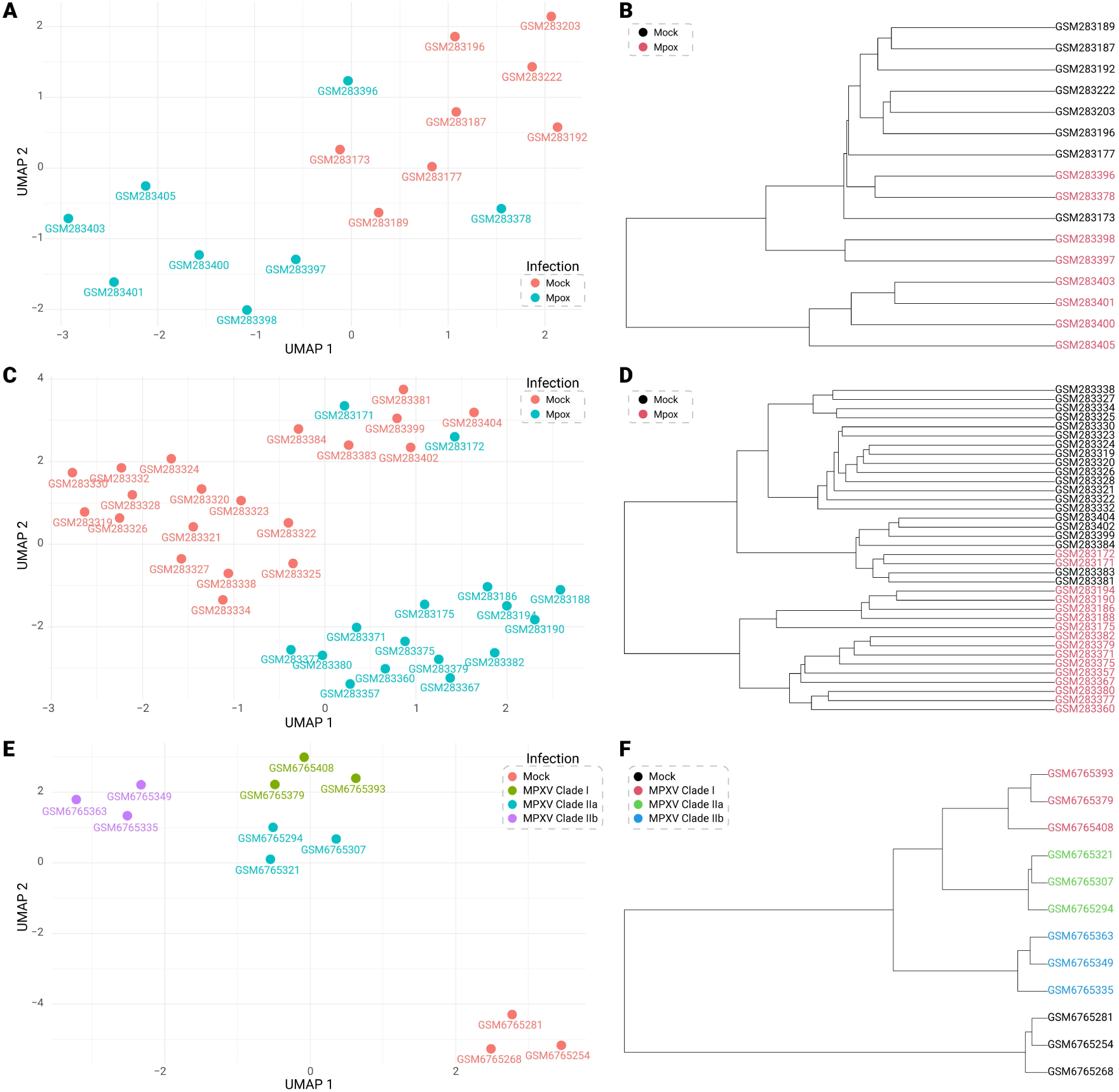
Sample distribution patterns and outlier identification through dimensionality reduction (UMAP, t-SNE) and phylogenetic analysis. (A, C, and E) Clusters among samples identified from UMAPs. X-axis represents the first component (C1), which captures the highest variation in gene expression while Y-axis represents the second component (C2) which delineates the second most variation in gene expression across the datasets. Each circle indicates a sample while varying colors indicating different treatments on the samples. (B, D, F) Phylogenetic trees visualized between the classes of samples to pinpoint the outlier samples.

#### Differential Gene Expression Analysis, Annotation, and Identification of DEGs

DGE analysis compared cell-line specific MPXV-infected samples to mock controls. In microarray data, 5,520 significant (padj < 0.05) annotated genes were evident in fibroblasts (4MPXV vs 8Mock), while 3,548 in monocytes (14 MPXV vs. 20 mock). Consequently, 922 up- and 1,849 down-regulated genes in fibroblasts (**Figure 6A**), and 590 up- and 277 down-regulated genes in monocytes (**Figure 6B**) were identified. Analyzing RNA-Seq data, substantial significant genes and DEGs were found in keratinocytes. After annotating significant Ensembl IDs, the following keratinocytes data were shown: (i) Clade I vs. mock: 2,631 upregulated and 2,212 downregulated (**Figure 6C**), (ii) Clade IIa vs. mock: 3,108 upregulated, 2,735 downregulated (**Figure 6D**), and (iii) Clade IIb vs. mock: 2,167 upregulated, 2,156 downregulated (**Figure 6E**).

**Figure 6.**
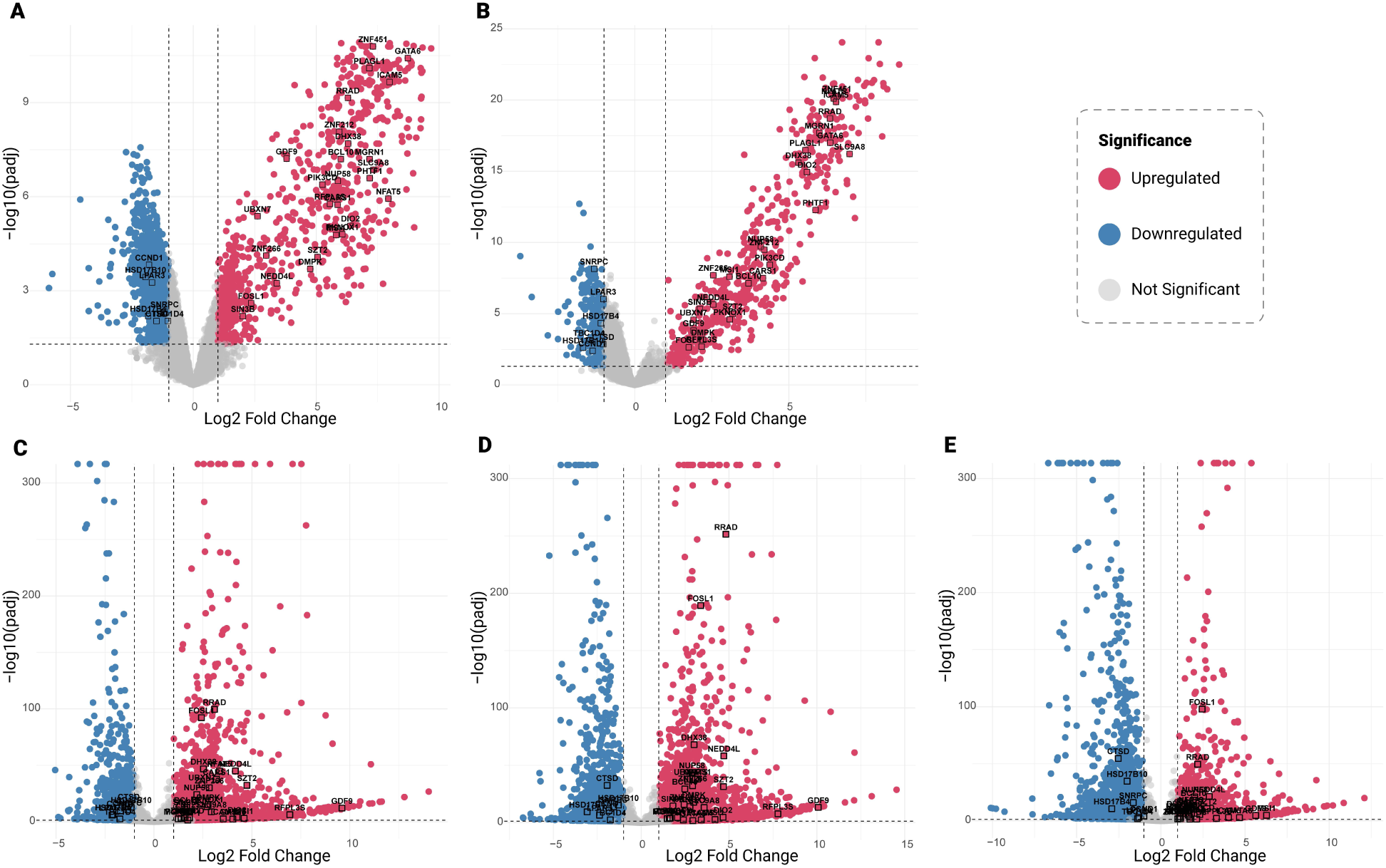
Upregulated and downregulated genes. X-axis denotes the log2 fold change (LFC) in gene expression. Positive values indicate upregulation whereas negative values indicate downregulation. Y-axis shows the negative log-transformed adjusted P-value. The red circles and the blue circles point out upregulated genes and downregulated genes, respectively. The horizontal line represents the threshold value of FDR smaller than 0.05, ascertaining the significance of the genes. However, the vertical lines represent the range of LFC less than −1 and greater than +1, nominating the significant genes as differentially expressed genes.

#### Shared DEGs and Conserved Expression Patterns Across Cell Types and Clades

Comparative analysis of DEGs revealed a conserved transcriptional response across MPXV clades (I, IIa, IIb) and cell types (fibroblasts, monocytes, and keratinocytes). Venn diagrams (**Figure 7A-B**) identified 27 up- and 7 downregulated genes common to all clades’ infection. Heatmap visualization (**Figure 7C**) of these 34 shared DEGs further delineated clade- and cell type-specific expression variability. This conserved signature underscores key genes are pivotal to MPXV pathogenesis, irrespective of viral lineage or host cell type.

**Figure 7.**
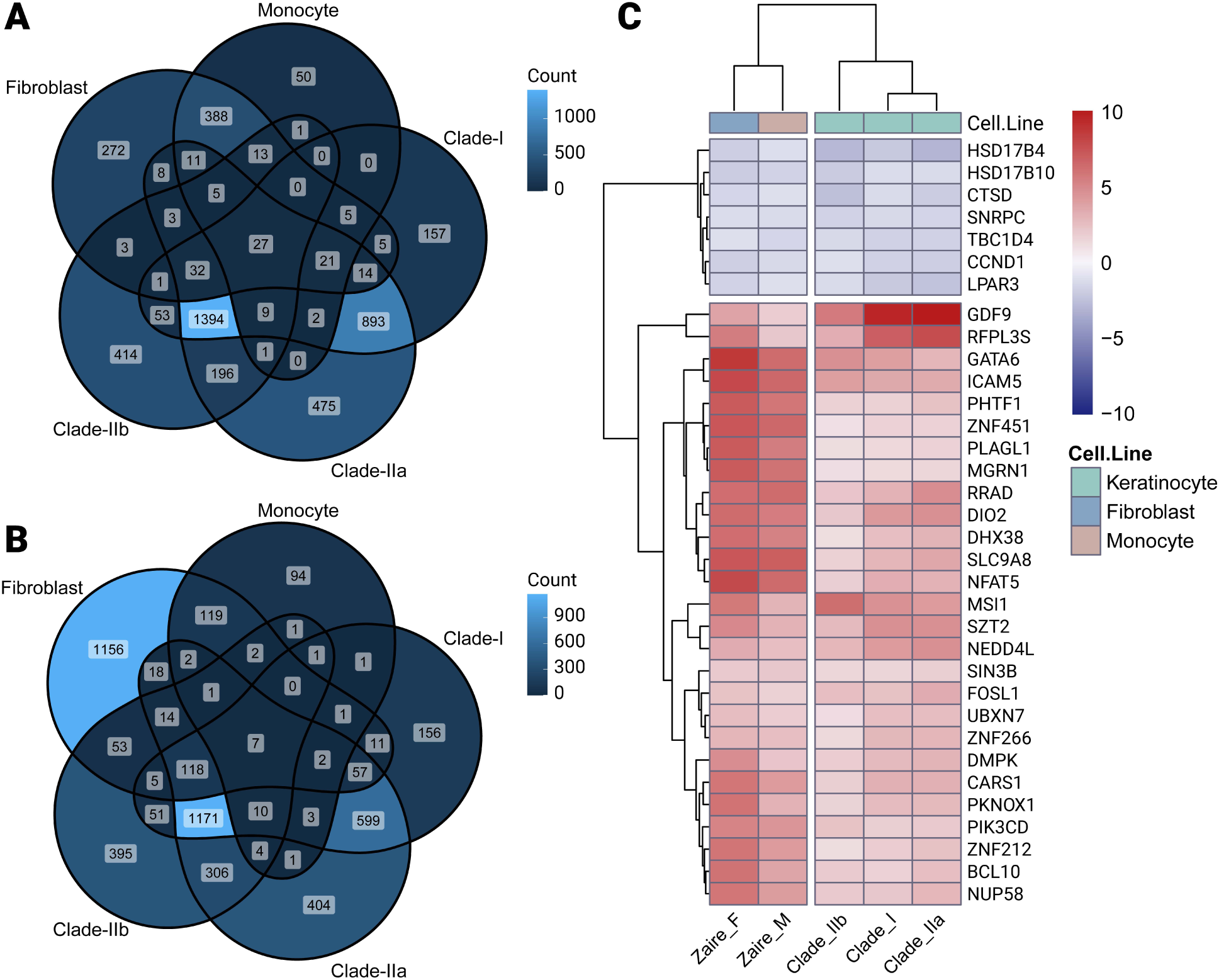
Comparative analysis and Heatmap. Venn diagrams depict the commonness and intersection in upregulated (A) and downregulated (B) genes. The heatmap (C) displays the expression patterns of the 34 shared DEGs based on log fold-change (LFC) values across different clades.

#### Integration of Machine Learning Framework

To enhance the reliability of biomarker discovery, automated machine learning (ML) algorithms were employed to evaluate the discriminatory power of identified DEGs in distinguishing MPXV-infected samples from controls. From comparative transcriptomic analyses, 34 DEGs conserved across all MPXV clades (I, IIa, IIb) were selected as initial features.

Primary data was constructed by merging normalized RNA-Seq (24 samples) and microarray datasets (124 samples), yielding 148 samples. Dimensionality reduction (PCA, t-SNE, UMAP), post batch effect correction and Z-score standardization, visualized the distribution of 148 samples based on the 34-feature expression profile (**Figure 8A-C**), while correlation analysis (**Figure 9**) assessed interdependencies among features. Feature selection refined the 34 DEGs to 17 non-redundant biomarkers using recursive feature elimination with random forests (RF-RFE). Using these 17 features, a reduced dataset was structured with the target classification (dependent variable).

**Figure 8.**
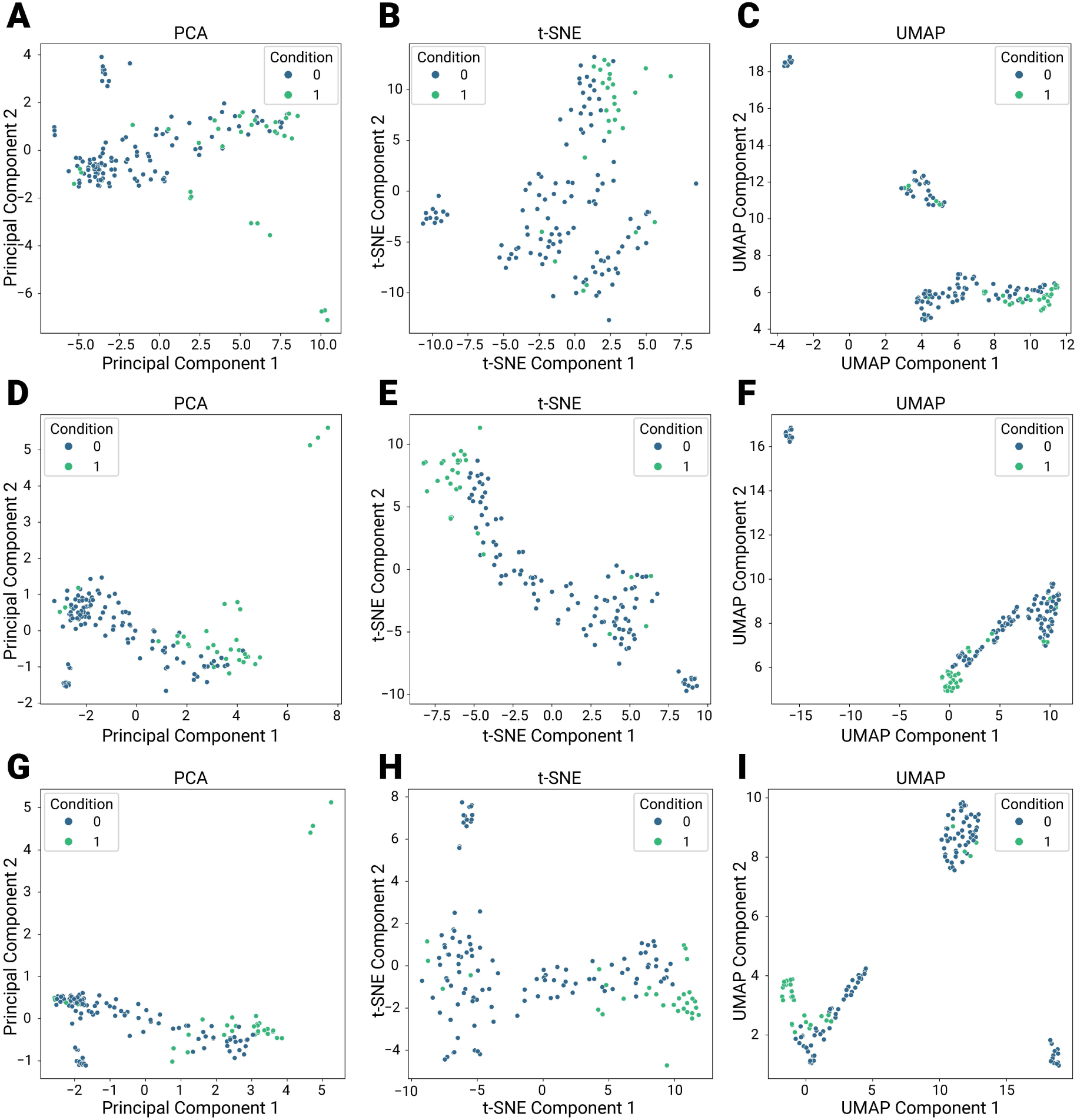
Assessment of the classification capabilities of features to cluster among the samples according to dimension reduction. The sample distributions are depicted using PCA (A), t-SNE (B), and UMAP (C) based on the 34 intersecting genes across infection of all clades of the Mpox virus. Similarly, distributions of samples are illustrated utilizing PCA (D), t-SNE (E), and UMAP (F) according to the top 10 genes ranked by importance score. Finally, discriminative power of the six genes, selected for the final model, is highlighted through PCA (G), t-SNE (H), and UMAP(I), plotted based on their expression levels. In the plots, each circle represents a sample while “1” represents Mpox samples, and “0” represents samples infected by other pathogens.

**Figure 9.**
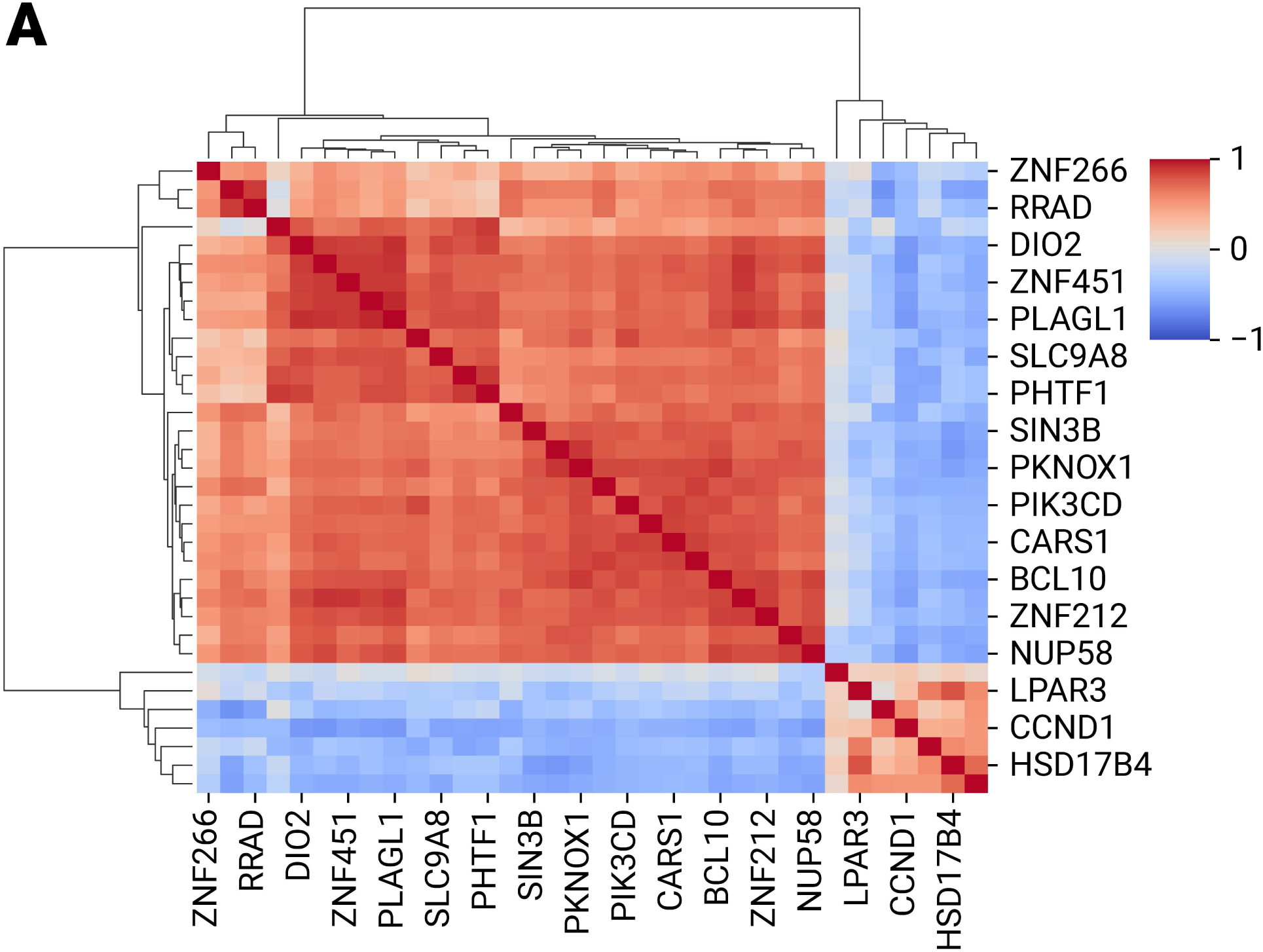
Relationships among the features shared to infection of all MPXV clades. The correlation plot depicts the interactions among the 34 features common to all clade infections. A correlation value of +1 indicates the highest positive correlation, while −1 represents the strongest negative correlation between two features. Positive correlations are represented by red, whereas blue signifies negative correlations.

#### Model Development and Benchmarking

By combining the most important features associated with MPXV infection into a Composite metric (disease indicator), a machine learning model, **markerMPXV,** was built. Twelve classification algorithms were rigorously tested iteratively. Using nested cross-validation and hyperparameter tuning, the Extra Trees (ET) classifier emerged as the top performer, achieving an accuracy of 0.95, AUROC of 0.97, AUPRC of 0.94, and F1 score of 0.90 (**Figure 10A-B**). Benchmarking results (mean ± std across outer folds) for all models are determined.

**Figure 10.**
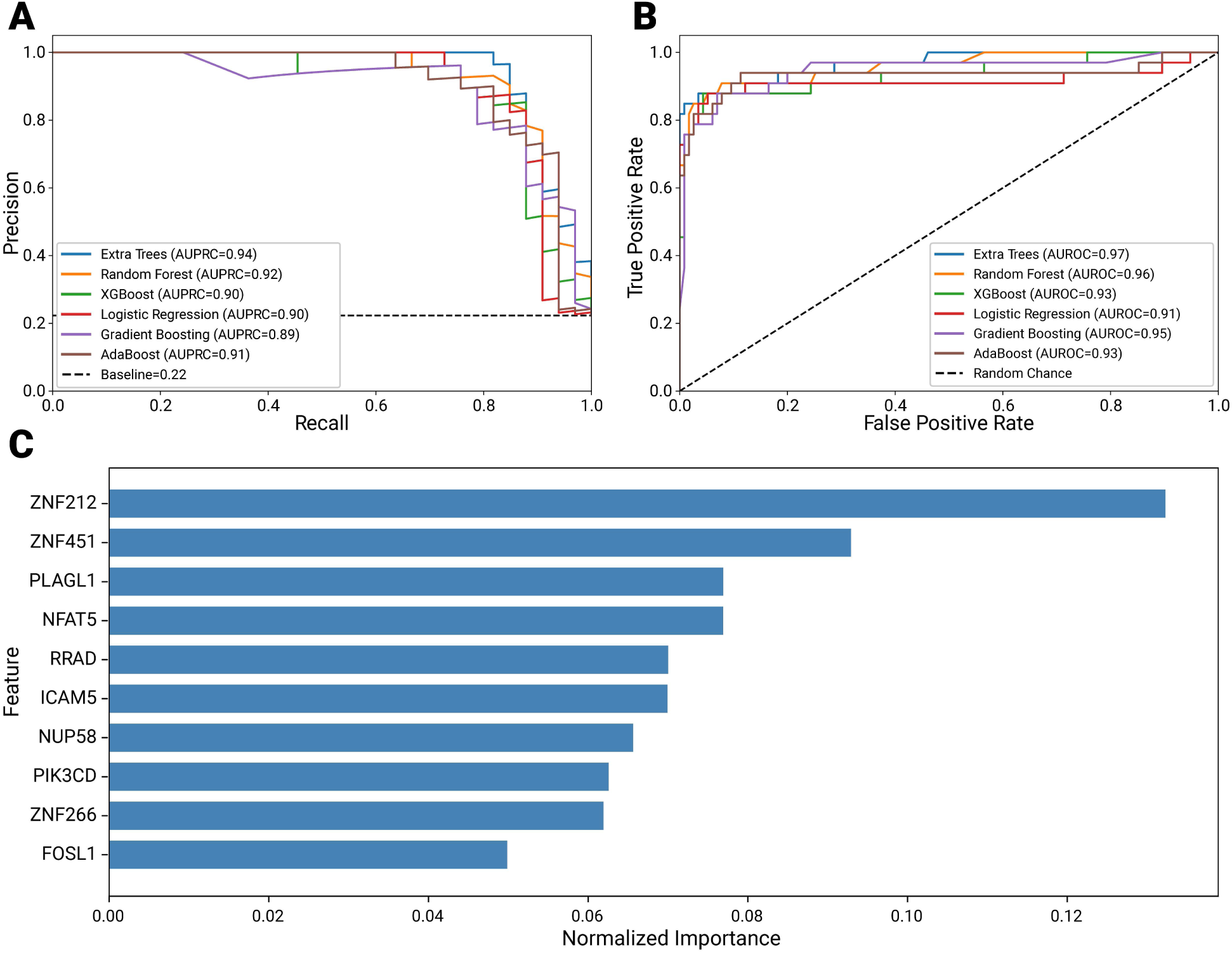
The benchmarking experiment identifies the best-fitting classifier for the dataset. (A) The AUPRC plot, also known as the precision-recall plot, illustrates the performance of six classifiers based on nested cross-validation. (B) The AUROC plot represents the accuracy of the classifiers to build models. (C) Highlights the ranking of the top 10 features based on their importance scores.

#### Feature Importance analysis and Single-Gene Models Facilitated Biomarker Prioritization by Ranking

The ET model ranked top 10 features by importance (**Figure 10C**). Single-gene Models ranked *RRAD* as the top among 10 biomarkers selected by feature importance. *RRAD* achieved robust performance (AUROC: 0.90; AUPRC: 0.85; F1: 0.76; accuracy: 0.91), demonstrating strong discriminative power between Mpox-infected and control samples (**Figure 11A-B**).

**Figure 11.**
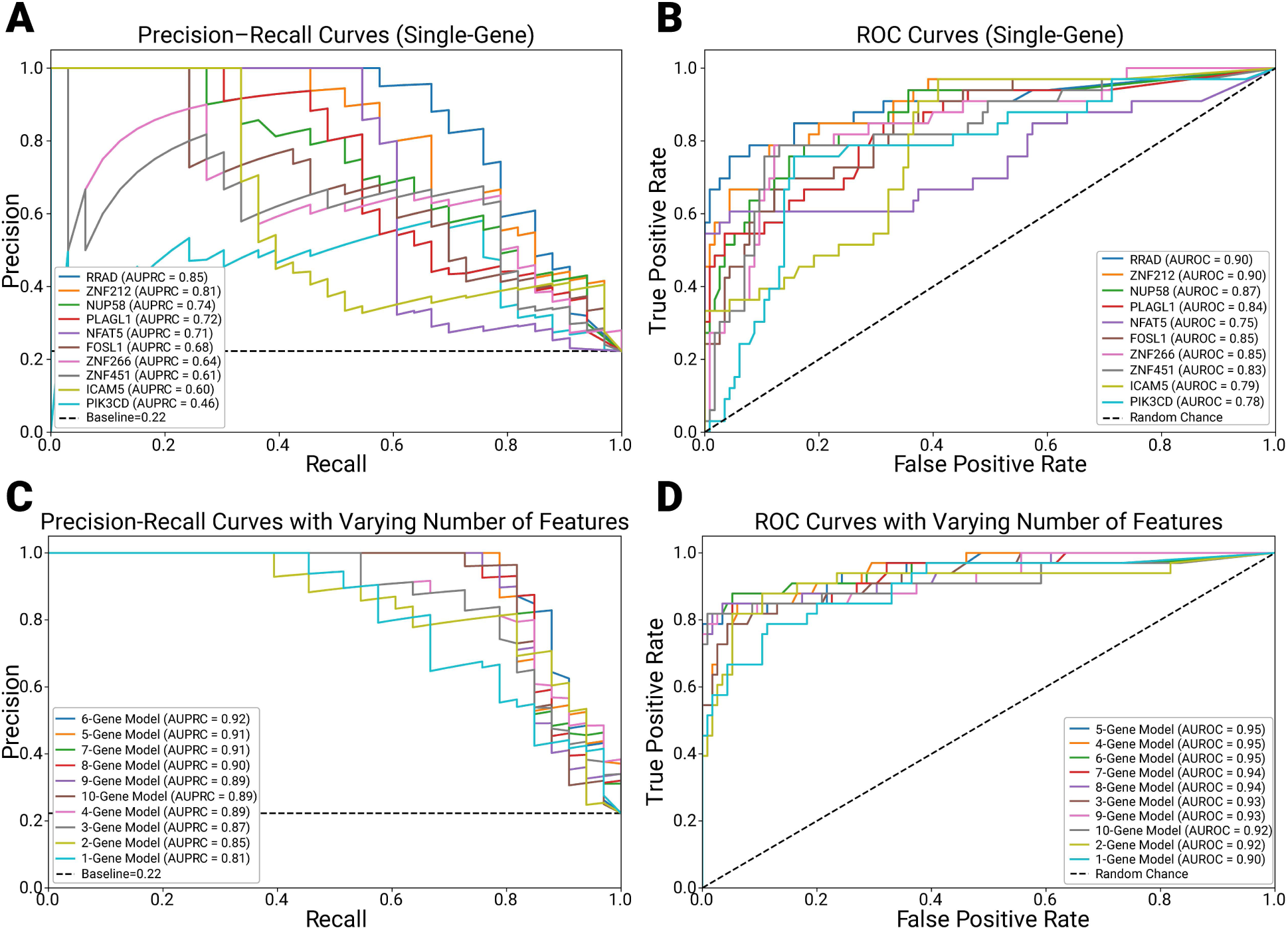
Evaluation of single- and multi-gene models. (A, B) The AUPRC and AUROC plots depict the performance and class-distinguishing ability of individual features. (C, D) illustrate the optimal combination of features for model building, determined by the ranking of performance (PR) and AUC scores. The evaluation ranking is based on AUPRC and AUROC values.

#### Feature Ablation and Optimal Multi-Gene Panel

A feature ablation study revealed that combining six genes (*ZNF212, ZNF451, PLAGL1, NFAT5, ICAM5, RRAD*) surpassed single-gene models, achieving superior performance (AUROC: 0.95; AUPRC: 0.92; F1: 0.84; accuracy: 0.94) (**Figure 11 C-D**). This six-gene panel was selected for final model refinement.

#### Final Model Performance and Validation

Consequently, the predictive model was built and finalized using the expression of the six genes (*ZNF212, ZNF451, PLAGL1, NFAT5, ICAM5, RRAD*), achieving an accuracy of 0.94 on train data (AUPRC: 0.91, AUROC: 0.93, F1: 0.84) and 0.93 on test data (AUPRC 0.91, AUROC 0.96, F1 0.83) (**Figure 12A-B**). The confusion matrices of train and test data for the finalized models are shown in **Figure 12C-D**. These results underscore the model’s reliability in diagnosing Mpox infection and its potential for clinical translation.

**Figure 12.**
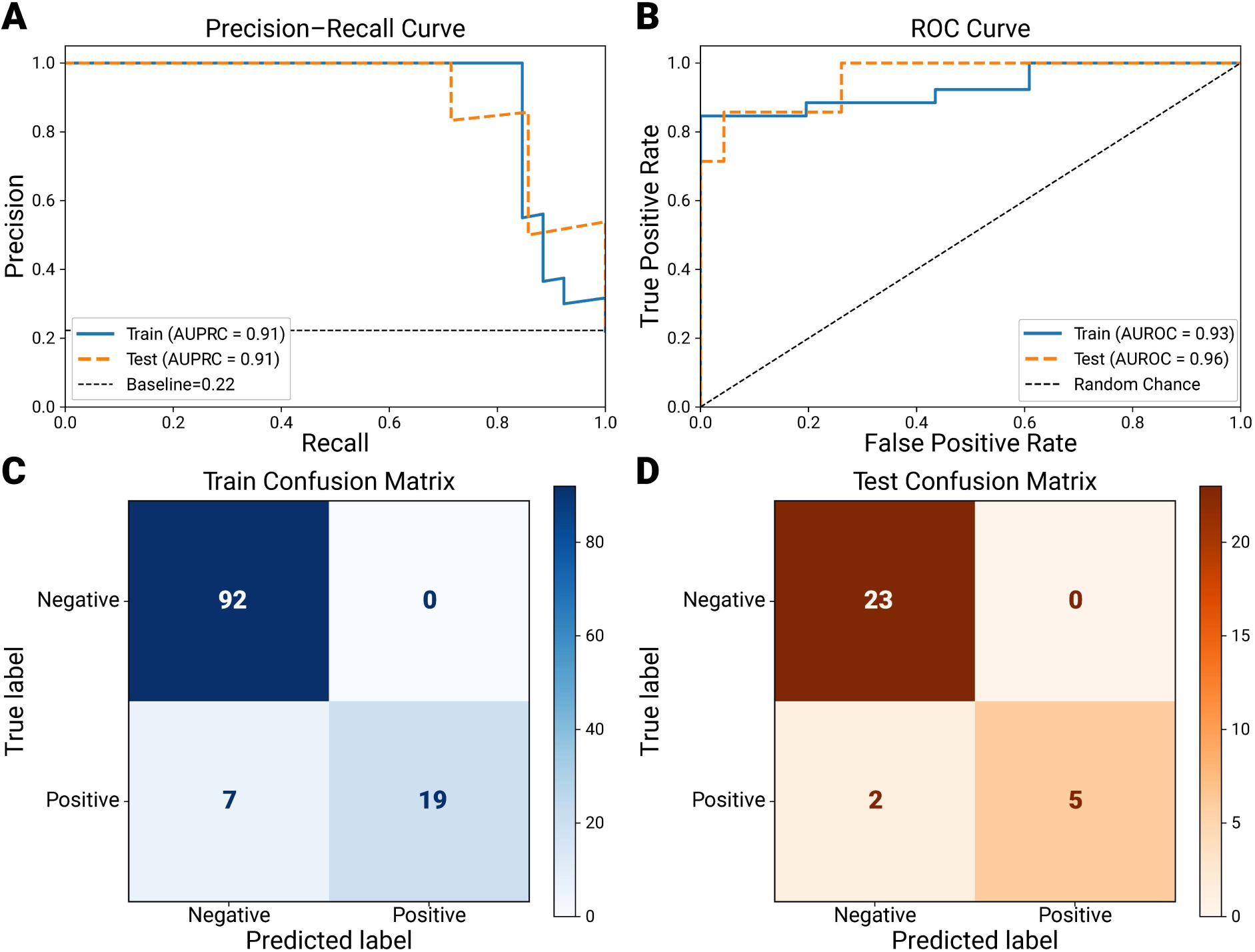
Performances of the Final Model X. (A, B) Model evaluation scores of AUPRC and AUROC, representing the performance of both training and testing of the final model. (C) Prediction outcomes on the test (C) and train (D) data are presented using the confusion matrix.

**Figure 13.**
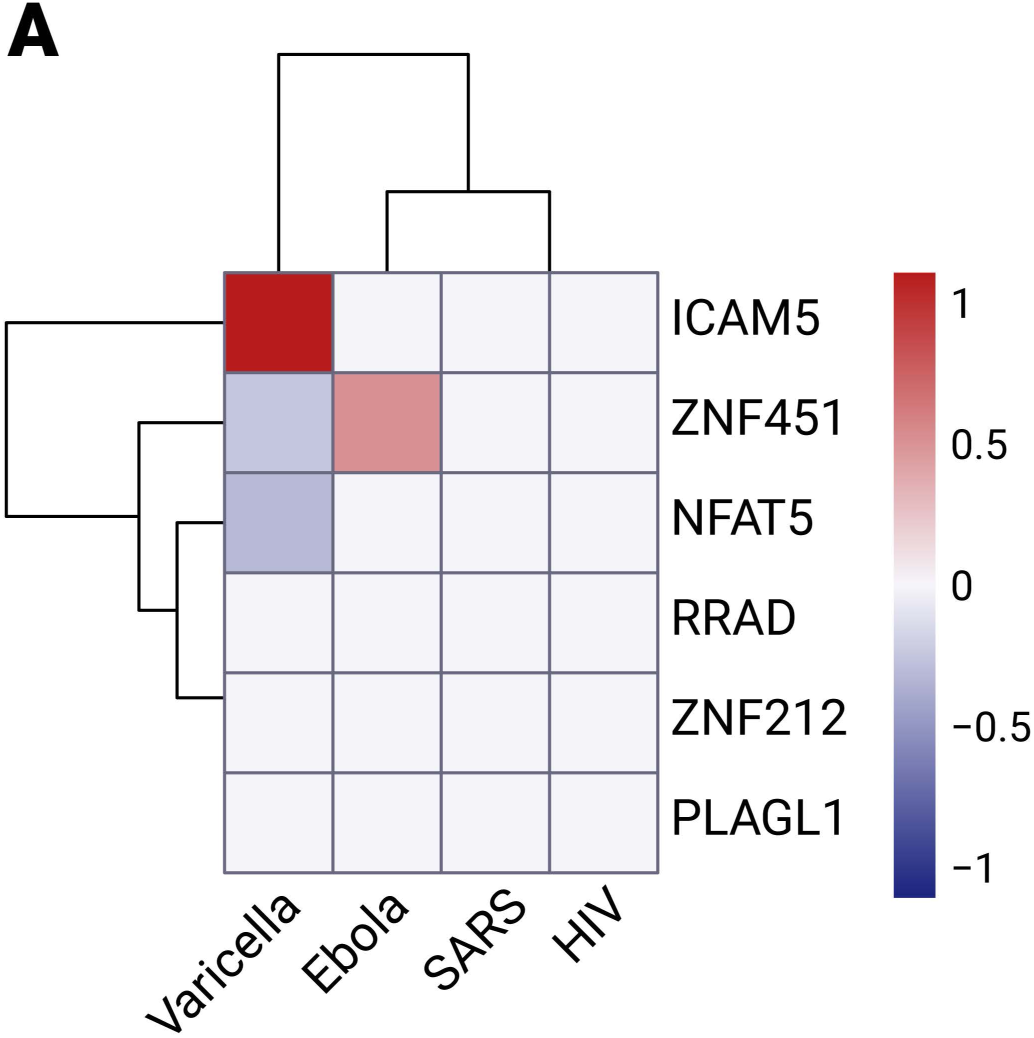
Expression level of selected biomarkers in other virus infection in human. The heatmap highlighting the scaled expression levels of six selected genes (ICAM5, ZNF451, NFAT5, RRAD, ZNF212, and PLAGL1) across four viral infection conditions: Varicella, Ebola, SARS, and HIV. The intensity of the colors reflects LFC values, where red indicates upregulation, blue indicates downregulation, and gray indicates no significance. Cladogram was applied to both genes and infection conditions to comprehend relationships pattern

#### Network, Functional, and Enrichment Analysis

To elucidate the functional roles of the six predicted biomarkers in MPXV infection, omicML mapped their interaction networks using the STRING database. 3,880 neighbor genes interacting with the six-gene panel (*ZNF212, ZNF451, PLAGL1, NFAT5, ICAM5, RRAD*) were identified. Each of the six genes were also analyzed individually and top 20 interacting neighbor genes’ network was plotted. Genes overlapping between the neighbor network and DEG lists were identified with enrichment analysis revealing their involvement in key biological processes and molecular functions.

#### Uniqueness of the Six-Gene Model as MPXV Biomarkers

To validate the hallmark signatures of Mpox infection of identified six-gene model including *RRAD*, transcriptomic datasets (GSE141932 [27], GSE157103 [28], GSE184320 [29], and GSE11234 [25]) of other potential viruses including varicella, ebola, HIV, and SARS-Cov-2 have been analyzed via omicML. Only ICAM5 (varicella) and ZNF451 (Ebola) exhibited marginal upregulation near significance thresholds, while other genes showed no differential expression. Interestingly, none of the six genes met statistical significance (padj < 0.05, |LFC| > 1) in smallpox-related varicella-zoster, SARS-CoV-2, HIV, or Ebola infections, confirming their specificity to MPXV. This lack of cross-viral relevance underscores the panel’s uniqueness as a robust MPXV signature.

## Discussions

The advent of high-throughput omics data and biomarker discovery techniques has resulted in a fragmented and specialized ecology of isolated platforms, rendering end-to-end analysis laborious. In practice, researchers frequently need to integrate disparate software for each task (e.g., DGE analysis in R/Python, annotation in external databases, GO pathway analysis in another tool) via manual file transfers and custom scripting, resulting in inefficiencies, errors, and reproducibility issues for non-programmers [30]. Most conventional pipelines (e.g., DESeq2, edgeR, limma) and annotation services (biomaRt, DAVID) are available solely as code libraries or standalone web applications, whereas point-and-click platforms (DEBrowser, GenePattern, GEO2R) generally cater to only a limited range of tasks, neglecting advanced procedures such as cross-study meta-analyses, batch correction, dataset integration, or machine learning (ML) investigations. Consequently, even standard procedures like quality control, normalization, and batch correction necessitate bioinformatics assistance, thereby constraining scalability and impeding workflow efficiency.

Moreover, workflows that use machine learning are typically absent. Traditional biomarker investigations often culminate with the compilation of a list of differentially expressed candidates, often evaluating them individually. Few tools offer built-in ML pipelines (feature selection, model training, benchmarking) to rigorously evaluate candidate biomarkers. However, Leclercq et al. indicated that current auto-ML systems are either inadequately designed for biological datasets or excessively complex for individuals lacking expertise in machine learning to utilize effectively [31]. In addition, the majority of machine learning technologies are inaccessible to biologists due to their support for a restricted array of algorithms, the necessity for manual hyperparameter optimization, or the assumption of programming proficiency [31]. Finally, inflexible, hard-coded pipelines lack the modularity and graphical interfaces necessary for seamless adaptation across various omics modalities or research strategies. The discipline presently needs a unified, adaptable platform that seamlessly integrates (1) DGE analysis, (2) annotation, (3) cross-study comparison, (4) ML-based validation, and (5) functional enrichment in an automated, user-friendly manner. Bench scientists encounter obstacles at every stage of biomarker discovery [30,32]. We introduce omicML, specifically engineered to address these deficiencies by offering a comprehensive, graphical platform for transcriptome biomarker identification.

All fundamental preprocessing and differential expression procedures are consolidated into a single platform, eliminating the necessity for users to transfer data across programs. Investigators can define experimental groups with minimal clicks and promptly obtain differentially expressed genes (DEGs) along with corresponding graphs. Integrated annotation (ID conversion, pathway mapping) and comparative modules (e.g., Venn overlaps across conditions) obviate the need for manual scripting or file transfers. The new approach eliminates the “silos” of disparate technologies, allowing for seamless transitions of outputs from data extraction to annotation to comparative modules. This immediately tackles the fragmentation problem identified in the literature.

omicML incorporates an extensive, machine learning-driven validation suite to overcome the shortcomings of traditional biomarker procedures. The platform transcends conventional methods that only identify statistical connections by automating feature selection, model benchmarking using nested cross-validation, and conducting ablation studies to thoroughly evaluate biomarker stability and significance. omicML employs tools such as BioDiscML for comprehensive searches and cross-validated classifiers, ensuring that biomarker candidates are evaluated using advanced methodologies without necessitating user coding [31].

provided as a graphical, no-code interface designed for anyone without a background in bioinformatics. Like BIOMEX, which illustrated the use of an interactive multi-omics platform for laboratory researchers, the new program offers menus and wizards in lieu of command lines. Non-experts may upload their data, configure parameters, and examine outcomes presented as publication-quality graphs and tables. The technology automates laborious activities such as batch correction and file merging, guaranteeing reproducibility without manual involvement. Bench scientists can go from raw data to validated biomarker panels solely within the GUI, eliminating the necessity for Python or R coding. This modular approach guarantees flexibility and reproducibility, allowing pipelines to be re-executed or modified for new datasets.

In our case study, we conducted a comparative transcriptomic analysis to identify DEGs and predictive biomarkers across multiple MPXV clades’ infection because of limited therapeutic options and the growing threat of a broader pandemic of monkey pox viruses. Using keratinocytes, dermal fibroblast, and monocyte cell-types infected with various MPXV clades, we found a higher number of DEGs in skin-derived cells compared to monocytes, a finding consistent with prior observations of increased viral load in keratinocytes **Figure 5 (A-E)**. Notably, recent clades appeared to elicit broader gene dysregulation compared to the older Zaire strain, with 34 DEGs (27 upregulated, 7 downregulated) consistently expressed across all three cell types irrespective of clade, suggesting their potential relevance in distinguishing MPXV pathogenesis **Figure 6 (A & B)**. To evaluate the diagnostic efficacy of these DEGs, we implemented machine learning models that classified MPXV-infected versus control samples using integrated RNA-Seq and microarray data. After batch effect correction, feature selection and benchmarking experiment, the Extra Trees Classifier uncovered *RRAD* as the most potent single-gene biomarker (AUROC: 0.90; AUPRC: 0.85; F1: 0.76; accuracy: 0.91). Furthermore, a six-gene panel (*ZNF212, ZNF451, PLAGL1, NFAT5, ICAM5, RRAD*) exhibited superior classification performance (AUROC: 0.95; AUPRC: 0.92; F1: 0.84; accuracy: 0.94), underscoring its utility for robust biomarker-based mpox detection. The identified biomarkers are pivotal in orchestrating host responses to MPXV infection, interacting with both upregulated and downregulated genes. Among the six key genes, five regulate critical cellular processes: *ZNF212, ZNF451, NFAT5*, and *PLAGL1* govern gene expression and biological pathways, while *RRAD* modulates molecular functions. The sixth gene, *ICAM5*, is central to cellular adhesion. Collectively, these genes form an interconnected network influencing signal transduction and immune responses, highlighting their systemic role in host-pathogen interactions.

*RRAD* and *ICAM5* are central to immune evasion [33,34]. *RRAD* suppresses NF-κB signaling by binding to its p50/p65 heterodimer, blocking inflammatory protein synthesis and cytokine production [34,35]. This inhibition dampens immune activation, potentially aiding MPXV survival. Notably, *RRAD* overexpression is linked to oncogenesis in skin cells and glucose metabolism dysregulation, contributing to type II diabetes [36,37]. Similarly, *ICAM5*, a neuronal immune modulator, is upregulated in MPXV-infected cells, impairing phagocytosis and T-cell responses [33,38]. Its overexpression may suppress innate and adaptive immunity, enhancing viral persistence and disease severity.

In contrast, *NFAT5* and *ZNF451* activate immune defenses. *NFAT5* promotes immune cell survival, proliferation, and differentiation (e.g., macrophages, T-cells) while regulating NF-κB and Treg/Th cell pathways. However, its overexpression risks rheumatoid arthritis and tumor progression, and may stimulate viral replication [39,40]. *ZNF451* enhances immunity by inhibiting TGF-β signaling, which otherwise suppresses NK cells, T-cells, and antigen-presenting cells [39]. By countering TGF-β, *ZNF451* amplifies immune activation, though its role in MPXV-specific responses warrants further study.

*PLAGL1* governs apoptosis, cell cycle control, and TP53-mediated transcription. As a tumor suppressor, its overexpression regulates aberrant proliferation yet is paradoxically associated with oncogenesis [41,42]. In MPXV infection, *PLAGL1*-induced apoptosis may restrict viral dissemination, while its multiple functions in cancer underscore context-dependent effects on host-pathogen interactions.

### Limitations

While omicML currently provides a comprehensive GUI-driven pipeline for transcriptomics-based biomarker discovery, more sophisticated functionalities are yet to be integrated in next versions (omicML 2.0). omicML is presently tailored to bulk transcriptomic data and does not include network-based or clinical modeling modules. In practice, many biomarker studies rely on gene co-expression network analysis and survival modeling to uncover complex patterns and clinical relevance, so these capabilities are absent in the current version.

## Conclusions

omicML represents a novel end-to-end framework for biomarker discovery by integrating many analytical steps into a cohesive, user-friendly platform. Its graphical interface guides users from data upload to normalization, differential expression, annotation, and machine-learning evaluation, therefore obviating the necessity for complex coding. This integrated pipeline unifies predictive modelling and biomarker selection into a cohesive approach. omicML democratizes access to complicated analyses by offering a GUI-based approach, allowing physicians and bench researchers without required programming abilities to execute advanced transcriptomics procedures. By reducing technical obstacles and offering a comprehensive, cohesive toolkit, omicML is positioned to significantly influence translational bioinformatics and the advancement of clinically pertinent molecular diagnostics.

Besides, omicML addressed the urgent need for mpox biomarkers and identified a six-gene model (*ZNF212, ZNF451, PLAGL1, NFAT5, ICAM5,* and *RRAD*) achieving exceptional diagnostic accuracy (AUROC: 0.95; AUPRC: 0.92) out of 34 clade-independent DEGs. This demonstrates omicML’s capacity to bridge transcriptomic insights with ML-driven validation, accelerating biomarker discovery for emerging pathogens and beyond.

## Abbreviations

GUI: Graphical User Interface
DGE: Differential gene expression
DEGs: Differentially expressed genes
LFC: Log2 fold change
FDR: False Discovery Rate
Padj: P-adjusted Value
PCA: Principal Component Analysis
UMAP: Uniform Manifold Approximation and Projection
t-SNE: t-distributed stochastic neighbor embedding
ML: Machine Learning
LR: Logistic Regression
ET: Extra Trees
RF: Random Forest
XGB: XGBoost
GB: Gradient Boosting
AB: AdaBoost
ACC: Accuracy
BACC: Balanced Accuracy
PREC: Precision
REC: Recall
F1: F1 Score
AUROC: Area Under the Receiver Operating Curve
AUPRC: Area Under the Precision-Recall Curve
MCC: Matthews Correlation Coefficient
KAPPA: Cohen’s Kappa
LOGLOSS: Log Loss
Mpox: Monkey Pox
MPXV: Monkey Pox Virus
GEO: Gene Expression Omnibus

## Acknowledgments

To the straightness of Kilo-Road, the sublimeness of Gazi Kalur Tila, and the symphonious rainfall in SUST Campus.

## Author Contributions

**Joy Prokash Debnath:** Methodology, Software, Formal analysis, Data Curation, Visualization, Investigation, Validation, Writing – original draft

**Kabir Hossen:** Methodology, Software, Formal analysis, Data Curation, Visualization, Investigation, Validation, Writing – original draft

**Md. Sayeam Khandaker:** Methodology, Software, Formal analysis, Data Curation, Visualization, Investigation, Validation, Writing – original draft

**Shawon Majid:** Software, Data Curation, Methodology, Validation, Investigation

**Md Mehrajul Islam:** Software, Data Curation, Methodology, Visualization, Investigation

**Siam Arefin:** Software, Investigation

**Preonath Chondrow Dev:** Conceptualization, Project administration, Resources, Supervision, Writing – review and editing

**Saifuddin Sarkar:** Conceptualization, Project administration, Resources, Supervision, Writing – review and editing

**Tanvir Hossain:** Conceptualization, Project administration, Resources, Supervision, Writing – review and editing

## Data availability statement

The datasets analyzed in the current study are available in the GEO repository. GSE219036: https://www.ncbi.nlm.nih.gov/geo/query/acc.cgi?acc=GSE219036 GSE11234: https://www.ncbi.nlm.nih.gov/geo/query/acc.cgi?acc=GSE11234 GSE141932: https://www.ncbi.nlm.nih.gov/geo/query/acc.cgi?acc=GSE141932 GSE157103: https://www.ncbi.nlm.nih.gov/geo/query/acc.cgi?acc=GSE157103 GSE184320: https://www.ncbi.nlm.nih.gov/geo/query/acc.cgi?acc=GSE184320

## Declarations

## Ethics approval and consent to participate

Not applicable.

## Consent for publication

Not applicable.

